# High-Resolution In Silico Painting with Generative Models

**DOI:** 10.1101/2024.05.31.596710

**Authors:** Trang Le, Emma Lundberg

## Abstract

Label-free organelle prediction presents a longstanding challenge in cellular imaging, given the promise to to circumvent the numerous drawbacks associated with fluorescent microscopy, including its high costs, cytotoxicity, and time-consuming nature. Recent advancements in deep learning have introduced numerous effective algorithms, primarily deterministic, for predicting fluorescent patterns from transmitted light microscopy images. However, existing models frequently suffer from poor performance or are limited to specific datasets, image modalities, and magnifications, thus lacking a universal solution. In this paper, we present a simplified VQGAN training scheme that is easily adapted with different input/output channels for image-to-image translation tasks. We applied the algorithm to generate multi-channel organelle staining outputs from bright field inputs, equivalent to the popular Cell Painting assay. The same algorithm also participated and placed first in the ISBI 2024 Light My Cell challenge.

## I. Introduction

Fluorescence microscopy is a commonly used technique in cell biology. However, the use of fluorescent probes in combination with the strong light needed to illuminate the fluorophore can introduce phototoxic effects and perturb cellular functions [1]. These limitations hinder the utility of fluorescence microscopy for long-term live cell imaging studies. Conversely, label-free transmitted light (TL) microscopy offers a non-invasive alternative with reduced phototoxicity, yet it does not inherently provide the specific cellular insights offered by targeting molecules by fluorescence probes. These label-free images captured through TL microscopy are believed to embed much more information about the cell and subcellular patterns than readily discernible to the human eye. Deep neural networks, which have demonstrated incredible effectiveness in natural image tasks, have proven powerful to unlock this latent information in biological images.

Recent advancements in deep learning, particularly image-to-image translation, have enabled innovative methods to predict organelle signals from TL images, bypassing the limits of traditional fluorescence microscopy. However, most approaches are limited to specific datasets or imaging modalities. For example, DeepHCS+ [2] used convolutional neural networks (CNN) for multi-task prediction of different channels (Nucleus, Cytosol and Apoptosis). Another notable study about label-free prediction [3] used UNET [4] to predict five Cell-Paint channels from bright-field images. Both approaches incorporate adversarial training (discriminator). A more complex training dataset, including three imaging modalities, cell lines, and data generated in three labs, was presented in a study where a CNN autoencoder was trained to predict organelles from TL inputs [5]. Despite being diverse in modality and cell line, this training dataset was still acquired at the same pixel size (as designed for the study), which helped CNN based models to recognize and learn perceptive fields. A more generalizable approach is needed to make use of previous microscopy studies, either intended or not intended for this task, as the public dataset pool grows.

Overall, there is great interest to develop generalized computational tools that can provide molecular labels from TL image input, and to validate such models in the most comprehensive way possible to facilitate downstream applications. For these tools to be easily usable by biologists, they need to be robust across a wide range of acquisition protocols, irrespective of the size of the images, cell line, acquisition site, modality or instrument. In the light of this aim, ISBI2024 Light My Cell (LMC) challenge recently made public a large diverse and heterogeneous dataset, including 30+ studies from 23 data acquisition sites, three main TL imaging modalities, multiple imaging settings and cell lines. Additionally, the Joint Undertaking in Morphological Profiling (JUMP) Cell Painting Consortium also gathers and makes available a massive public CellPaint datasets [6]. These resources offer great opportunities to develop methodology aimed for greater generalizability across diverse biological specimens and imaging conditions. This paper outlines our first-place approach to the LMC challenge, using relatively light-weight generative models for image-to-image translation tasks. Our approach also surpassed metrics of the previously published label-free CellPaint [3] (albeit different dataset). The resulting models bring us a step closer to obtaining molecular readouts from non-invasive cellular imaging technology, offering profound opportunities for biological research.

## II. Method

### A. Datasets

#### CellPaint dataset

A subset of the JUMP pilot dataset (`cp0000`) [6], [7] was downloaded and preprocessed to test model capacity when all organelle channels are present. In specific, 4429 large field of views (FOVs) from unperturbed U2OS cells were each divided into 16 tiles. Each FOV contains 8 channels: 3 bright field (BF) and 5 organelle channels (DNA, ER, RNANucleoli, AGP and Mitochondria). After removing empty and low cell count tiles, the final JUMP dataset included 58237 tiles. This dataset was acquired with a widefield microscope at 20x objective. This dataset was split into train/validation/test at 80%/10%/10% proportion, stratified by plate.

#### LMC Challenge dataset

The challenge dataset includes 56700 previously unpublished images from 30 studies from different sites, containing TL channels and 4 organelles: Nucleus, Actin, Mitochondria and Tubulin. This dataset is very heterogeneous, including 3 TL imaging modalities (Bright Field, Phase Contrast, Differential Interference Contrast microscopy), 40x - 100x objective, pixel sizes from 65 to 110 nm, 11 different human and dog cell lines. Each study consists of one mode of TL (multiple z-stacks), 1-3 organelles stained in one cell line. In terms of organelle class, there is a great class imbalance, i.e. 2 orders of magnitude difference in the number of training data for Actin (27 images) compared to Nucleus (2533 images) (Figure 1A). Furthermore, among 2574 unique non-empty FOVs, each has 1-3 organelle channel combinations and there’s no FOV with all organelles, as presented in Figure 1B.

**Fig. 1.**
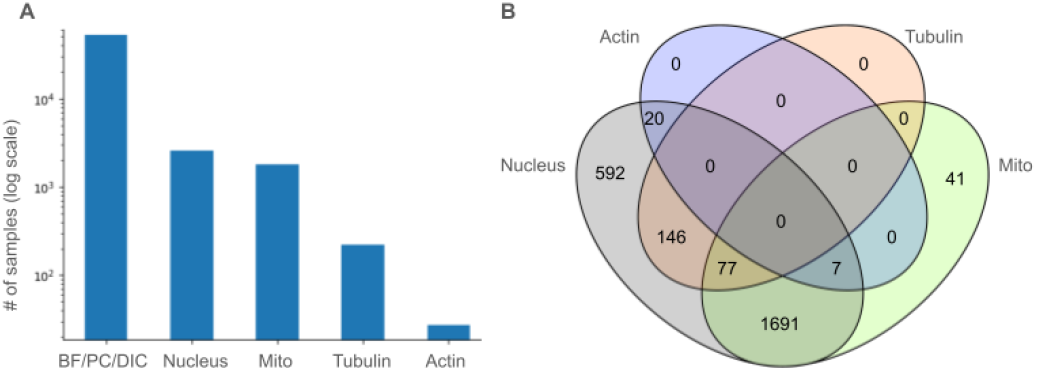
Challenge data distribution. A. Number of images from challenge dataset in each class: transmitted light modalities and 4 organelles. B. Venn diagram of number of field of views with certain channel combinations.

The LMC challenge dataset was split into train/validation at 95%/5%, stratified split by studies. For the competition, participants submitted 1 algorithm that would predict 4 organelle channels from each TL input. There is also a strict limit on computational resources that each algorithm can use, for real-world applicability and leveling the playing field. At test time, all participating algorithms were evaluated with a wide range of metrics for each organelle and 6 z-stack deviations. The held-out test set consists of 322 FOVs, including one additional acquisition site completely separated from the challenge dataset.

### B. Evaluation Metrics

A combination of metrics were used to evaluate all algorithm submissions in the LMC challenge, including Mean Absolute Error (MAE), Structural Similarity Index (SSIM), Pearson Correlation Coefficient (PCC), Euclidean and Cosine Distances (*E dist & C dist*). For the LMC challenge, for each input (1 TL z-stack), four organelle outputs were predicted. For Nucleus and Mitochondria, all five metrics were used. For the filamentous organelles Tubulin and Actin, only SSIM and PCC were used. A ranking for each organelle was determined by all metrics calculated from 0-to-5 deviations from the focus plane where specific organelle patterns were captured. Participants were ranked based on this 4 organelle x 6 deviation metrics matrix, with winners determined by the best average across all metrics. For the CellPaint dataset, all metrics were calculated for all output channels.

### C. Model architecture

The generative architectural framework we chose for this study is a simplified version of the VQGAN [8] (without taming transformer), selectively retaining crucial components to ensure good performance while maintaining simplicity and relatively low computational burden during training. The architecture (Figure 2) incorporates an encoder for data compression and a decoder for output prediction. Quantization is applied within the embedding space to promote a more generalized codebook. Notably, the decoder functions as a generator, synthesizing organelle channels based on the quantized vector. Moreover, a discriminator is incorporated into the prediction process to foster the generation of realistic output representations.

**Fig. 2.**
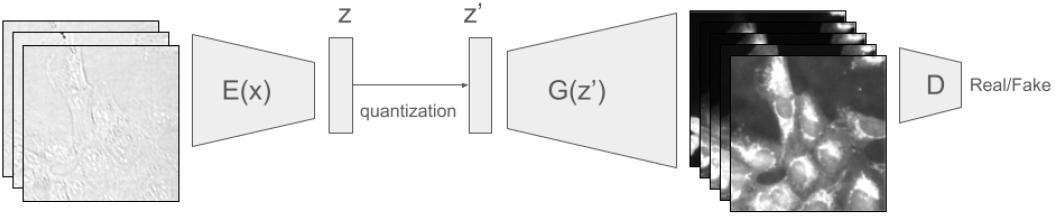
Model architecture of VQGAN with an encoder for data compression, a quantized embedding space for generalization, and a generator that generates organelle channels. A discriminator ensures realistic output representations.

In specific, we define two image domain X (transmitted light) and Y (organelle staining), the aim is to provide a transformation *f* : X →Y. The encoder, *E*, compresses the input data *x* ∈ *R*^*H×W ×C*^ into a latent representation 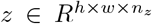. The latent representation *z* is quantized to *z*^*′*^ using a element-wise quantization q(·), which maps each spatial code *z*_*ij*_ to the nearest codebook entry:

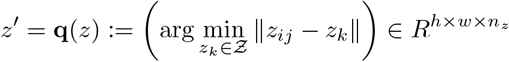

The codebook loss, *ℒ* _*codebook*_, ensures the quantized vectors *z*^*′*^ are close to the encoder output *z* while also encouraging the encoder output to move towards the codebook entries:

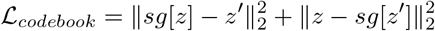

where *sg*[*·*] denotes the stop-gradient operation [9].

The decoder, *G*, reconstructs the high-resolution output 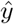 from the quantized latent representation *z*^*′*^. To facilitate high-resolution output, the decoder is larger than the encoder, with 1.5 times the number of trainable parameters.

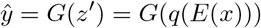

The pixel loss, *ℒ* _*pixel*_, measures the reconstruction error between the target *y* and the output 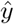:

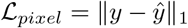

A discriminator, *D*, is incorporated into the prediction process to foster the generation of realistic output representations.

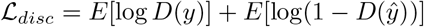

The overall training objective is to minimize the combined loss:

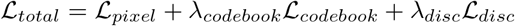

where *λ*_*codebook*_ and *λ*_*disc*_ are weighting factors.

Each VQGAN model has 58.1 million trainable parameters.

#### Adaptation for channels

The architecture easily adapts to various input and output channels for biological image data. Without a perceptual loss pretrained on natural images (lpips with vgg pretrained on Imagenet [10]), the model is not restricted to 3-channel inputs/outputs. In this study, it handles 3-channel inputs and produces outputs with 5 or *n* channels, offering flexibility f or d ifferent b iological imaging applications.

### D. Training Scheme

#### Preprocessing

To reduce batch effects and intensity range difference, each image was normalized between 2% and 99.8% pixel percentile. Simple augmentation techniques, including flipping along 2 principal axes, 90 rotations, and random cropping and resizing ranging from 70% to 95% of the image size, were employed to somewhat address the different resolutions of studies ranging from 20x to 100x objectives. After augmentation, each sample was resized to the final input size of 256x256 before feeding in the model.

#### Experiments

To check the performance of the specialized organelle models compared to the combined model, we train seven models on the same train/validation/test split for on the subset of JUMP pilot dataset described above. The experiments and performance is outlined in Table III and Appendix A. Each model was trained on 1 NVIDIA A100 GPU, with ADAM optimizer, learning rate of 4.0e-06, batch size of 20, and for 20 epochs (20h). For LMC, four organelle specific models were trained, each on 1 NVIDIA A100 GPU, with ADAM optimizer and learning rate of 2.0e-06 (for Actin), 4.0e-06 (for Nucleus), 1.0e-06 (for Mitochondria and Tubulin), and batch size of 16. Validation was performed every epoch for Tubulin and Actin, and every half epoch for Nucleus and Mitochondria. See more details about the submission in Appendix B.

## III. Result

### A. Organelle-specifc vs combined model performance

We compared the performance of organelle-specific models to a combined model by systematically varying the input and output configurations in the CellPaint dataset. First, we observed that more input (BF) information boosted model performance, as models with all 3 BF channels outperform single BF inputs for all five organelles. Second, with the same data split and computational resources, specialized models still outperformed one combined model, consistently from the worst (Mitochondria) to the best (DNA) performing organelle class (Table III). Image outputs from the combined model are visualized in Figure A.1.

### B. Model performance in LMC challenge

Our algorithm submission to the LMC challenge consists of 4 organelle specific models. Some sample images from the validation set are visualized in Figure A.2. On the final hidden test set, our method achieved the average ranking of 2.4, surpassing the 2nd solution by 1.0 point. Our solution placed first for 6/14 metrics across organelles. Notably, there doesn’t seem to be a great agreement among the “intensity” (PCC, MAE), structure (SSIM) or “texture” (*E dist & C dist*) metrics, as defined by the challenge. For example, our Nucleus model placed 1st for 3/5 metrics, but only 4th on the other 2 metrics (Table A.1). This disagreement is beneficial for a more comprehensive evaluation, as a combination of diverse metrics, each capturing different aspects of model performance and potentially offering conflicting perspectives, can account for various strengths and weaknesses of each channel.

## IV. Discussion

With advances in deep learning, label-free organelle prediction is becoming feasible. We present a simplified VQGAN training scheme, which can be applied flexibly to different biological image datasets, and tested this approach on two large and diverse TL-to-organelles datasets. This approach outperforms other solutions (UNET and GANs) in ISBI 2024 LMC challenge to predict organelle patterns through TL images. The winning solution includes individual models trained to predict specific organelles (specialized models). While the extreme class and study (output combination) imbalances inherent in the competition context posed great challenges for devising a universally optimized training scheme for all output types, a combined model whose encoder learnt common features from TL inputs would be much more efficient and might provide the generalizability that large models are after (for e.g. through regularization). However, experiments with the CellPaint dataset demonstrate that, with the same resources, specialized models still outperform one combined model. Moreover, in a low-data regime, specialized models can be trained with different hyperparameters and can maximize performance separately. Additionally, they allow for organelle-specific improvements, for e.g. when there’s more training data for just one certain class, without affecting other outputs, which might be preferable for many researchers.

**TABLE 1.** Performance Metrics for Different Image Channels and Model Inputs on CellPaint dataset.

| Image channel | Model output | Model input | MSE↓ | SSIM↑ | MAE↓ | PCC↑ | Edist↓ | Cdist↓ |
| --- | --- | --- | --- | --- | --- | --- | --- | --- |
| DNA | DNA | 3 BF channels | **207.45** | **0.92** | **5.82** | **0.97** | **3501.84** | **0.03** |
|  | 5 channels | 3 BF channels | 297.03 | 0.91 | 6.51 | 0.95 | 4221.56 | 0.04 |
|  |  | BF 1 | 475.57 | 0.88 | 8.22 | 0.94 | 5356.85 | 0.06 |
|  |  | BF 2 | 391.17 | 0.89 | 7.55 | 0.94 | 4855.18 | 0.05 |
|  |  | BF 3 | 833.86 | 0.85 | 10.59 | 0.85 | 6983.43 | 0.06 |
| ER | ER | 3 BF channels | **333.18** | **0.85** | **9.55** | **0.95** | **4473.44** | **0.03** |
|  | 5 channels | 3 BF channels | 483.46 | 0.80 | 11.72 | 0.92 | 5418.48 | 0.02 |
|  |  | BF 1 | 656.83 | 0.76 | 13.61 | 0.90 | 6349.34 | 0.03 |
|  |  | BF 2 | 682.02 | 0.76 | 13.88 | 0.89 | 6466.65 | 0.03 |
|  |  | BF 3 | 927.12 | 0.74 | 16.81 | 0.84 | 7553.60 | 0.04 |
| RNANucleoli | RNANucleoli | 3 BF channels | **244.95** | **0.90** | **8.66** | **0.97** | **3796.91** | **0.01** |
|  | 5 channels | 3 BF channels | 330.34 | 0.87 | 10.16 | 0.96 | 4444.67 | 0.02 |
|  |  | BF 1 | 502.32 | 0.82 | 12.65 | 0.94 | 5508.25 | 0.03 |
|  |  | BF 2 | 530.29 | 0.81 | 13.10 | 0.93 | 5670.64 | 0.03 |
|  |  | BF 3 | 791.99 | 0.72 | 16.78 | 0.89 | 6969.78 | 0.05 |
| AGP | AGP | 3 BF channels | **414.68** | **0.81** | **10.74** | **0.92** | **5035.28** | **0.03** |
|  | 5 channels | 3 BF channels | 525.75 | 0.78 | 12.36 | 0.90 | 5709.45 | 0.04 |
|  |  | BF 1 | 654.37 | 0.57 | 13.97 | 0.88 | 6378.49 | 0.06 |
|  |  | BF 2 | 667.41 | 0.74 | 14.23 | 0.88 | 6450.29 | 0.04 |
|  |  | BF 3 | 869.75 | 0.69 | 16.89 | 0.86 | 7359.61 | 0.07 |
| Mitochondria | Mitochondria | 3 BF channels | **683.10** | **0.70** | **18.74** | **0.87** | **6205.44** | **0.02** |
|  | 5 channels | 3 BF channels | 955.22 | 0.65 | 22.34 | 0.83 | 7361.92 | 0.03 |
|  |  | BF 1 | 1336.08 | 0.57 | 27.90 | 0.77 | 8782.18 | 0.03 |
|  |  | BF 2 | 1252.43 | 0.59 | 26.55 | 0.77 | 8528.63 | 0.07 |
|  |  | BF 3 | 1311.16 | 0.56 | 26.69 | 0.73 | 8852.29 | 0.04 |

## Software and Data

Code and documentation can be found at https://github.com/trangle1302/lmc_GenCellPaint JUMP CellPaint dataset are available from the Cell Painting Gallery on the Registry of Open Data on AWS https://registry.opendata.aws/cellpainting-gallery/. LMC Challenge dataset are available to download at: https://lightmycells.grand-challenge.org/database/.

## Acknowledgment

The authors would like to thank the JUMP-CP consortium and the organizers of ISBI2024 Light My Cell competition and all data generators for gathering and making the diverse datasets public.

## APPENDIX

**TABLE A.1.**
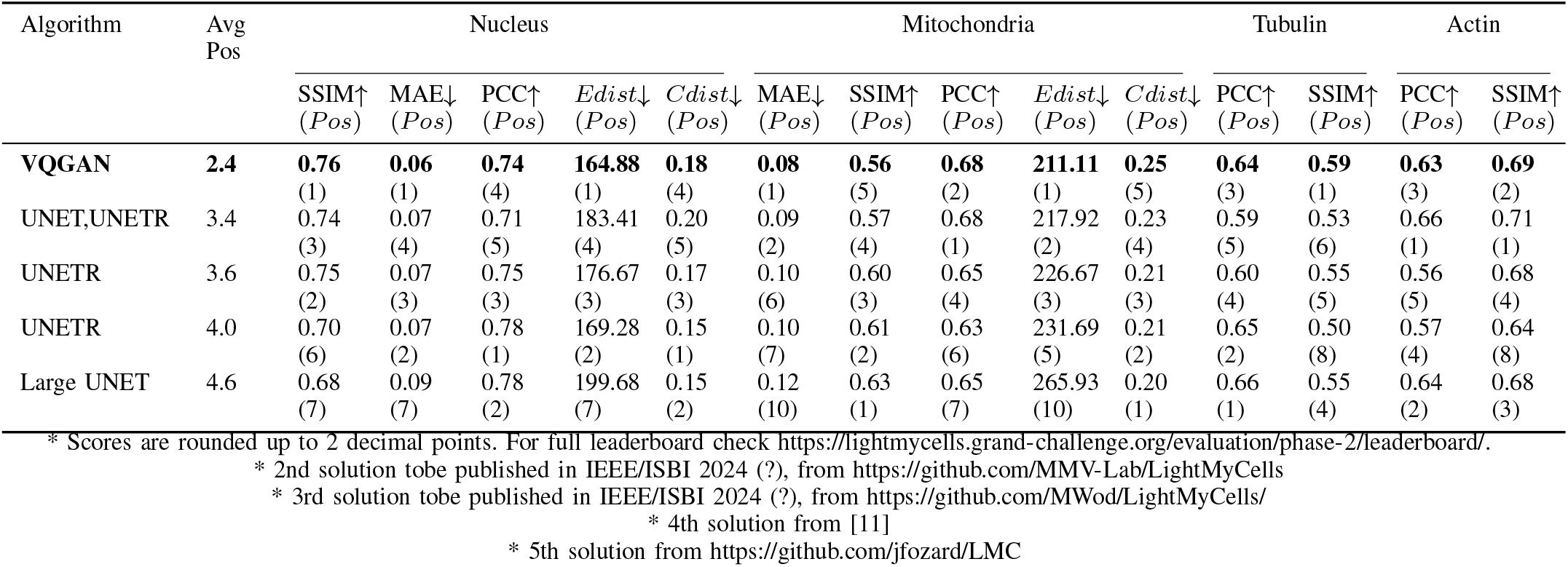
Performance Evaluation of Algorithms on phase 2 hidden test set of LMC Challenge

### A. etails for experiments on CellPainting

We trained 7 models with these combinations of inputs/outputs:

- 3 BF channels →5 organelle channels
- 1 BF channel (chosen from BF1, BF2, BF3) →5 organelle channels
- 5 organelle specific models: 3 BF channels →DNA | ER | RNANucleoli | AGP | Mitochondria

### B. Details for first place submission to LMC challenge

While some models just started to converge, the training time at submitted checkpoints varies for each model:

- Model for Nucleus was trained for 54 epochs, average train (and validation) time of 2h/epoch.
- Model for Mitochondria was trained for 55 epochs, average train (and validation) time of 2.2h/epoch.
- Model for Tubulin was trained for 309 epochs, average train (and validation) time of 24 min/epoch.
- Model for Actin was pre-trained for 24 epochs on JUMP dataset (70 min/epoch), then 122 epochs fine-tuning with average training time of 15 min/epoch. Pretraining/Finetuning details: CellPaint was used for pre-training for the Actin prediction model, specifically the middle bright field (ch07, BF2) and AGP (Actin, Golgi, Plasma membrane, ch02) channels. After the models learnt basic cell morphology from the background in transmitted light images, the training data was updated to employ 1/5th of the JUMP data from pretraining in addition to the LMC training set, with validation exclusively performed on the provided LMC validation set.

**Fig. A1.**
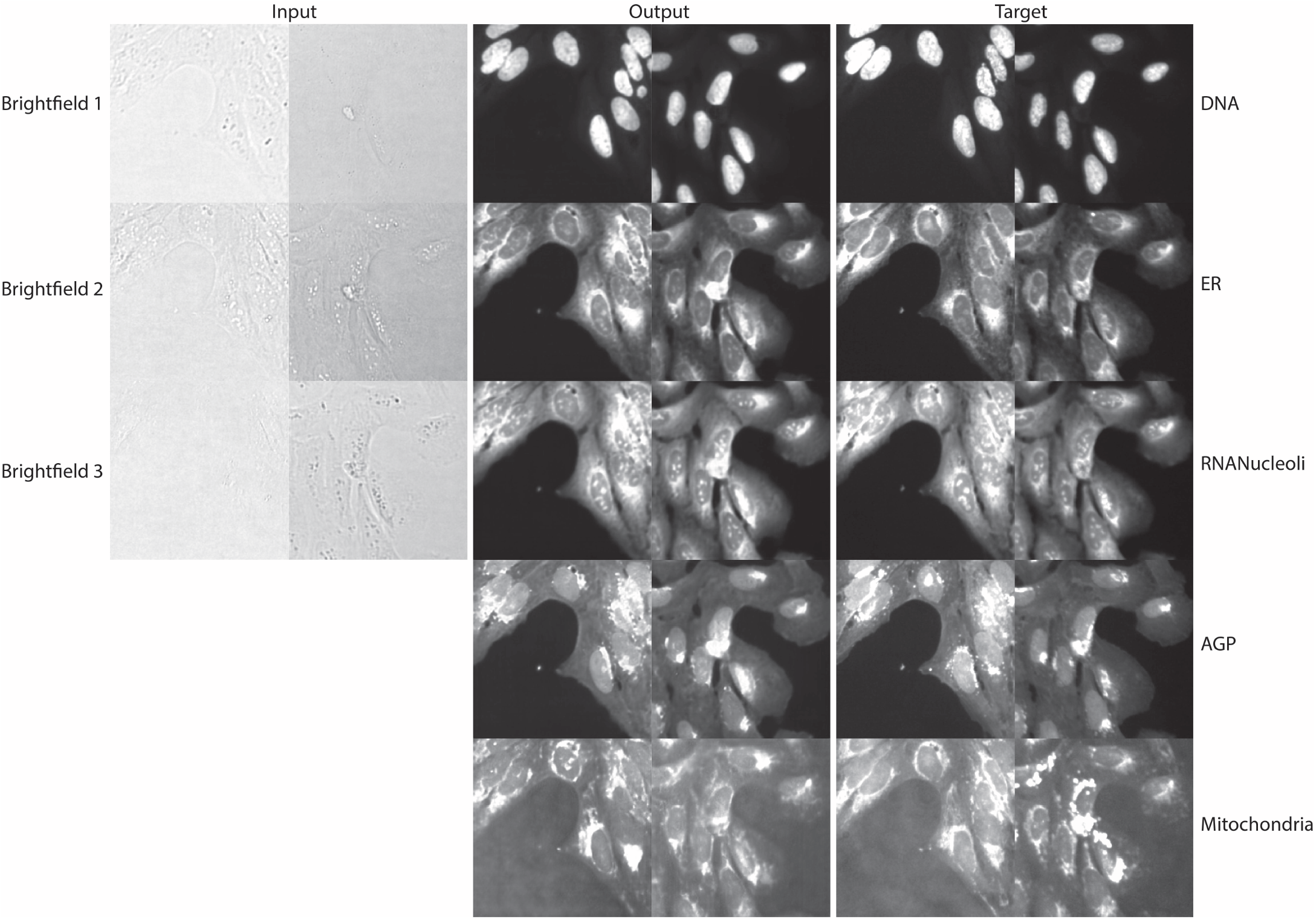
3 BF channel to 5 organelle model outputs on test images from JUMP CellPaint dataset, sampled at random. Inputs and Targets are rescaled to [2,99.8] percentile.

**Fig. A2.**
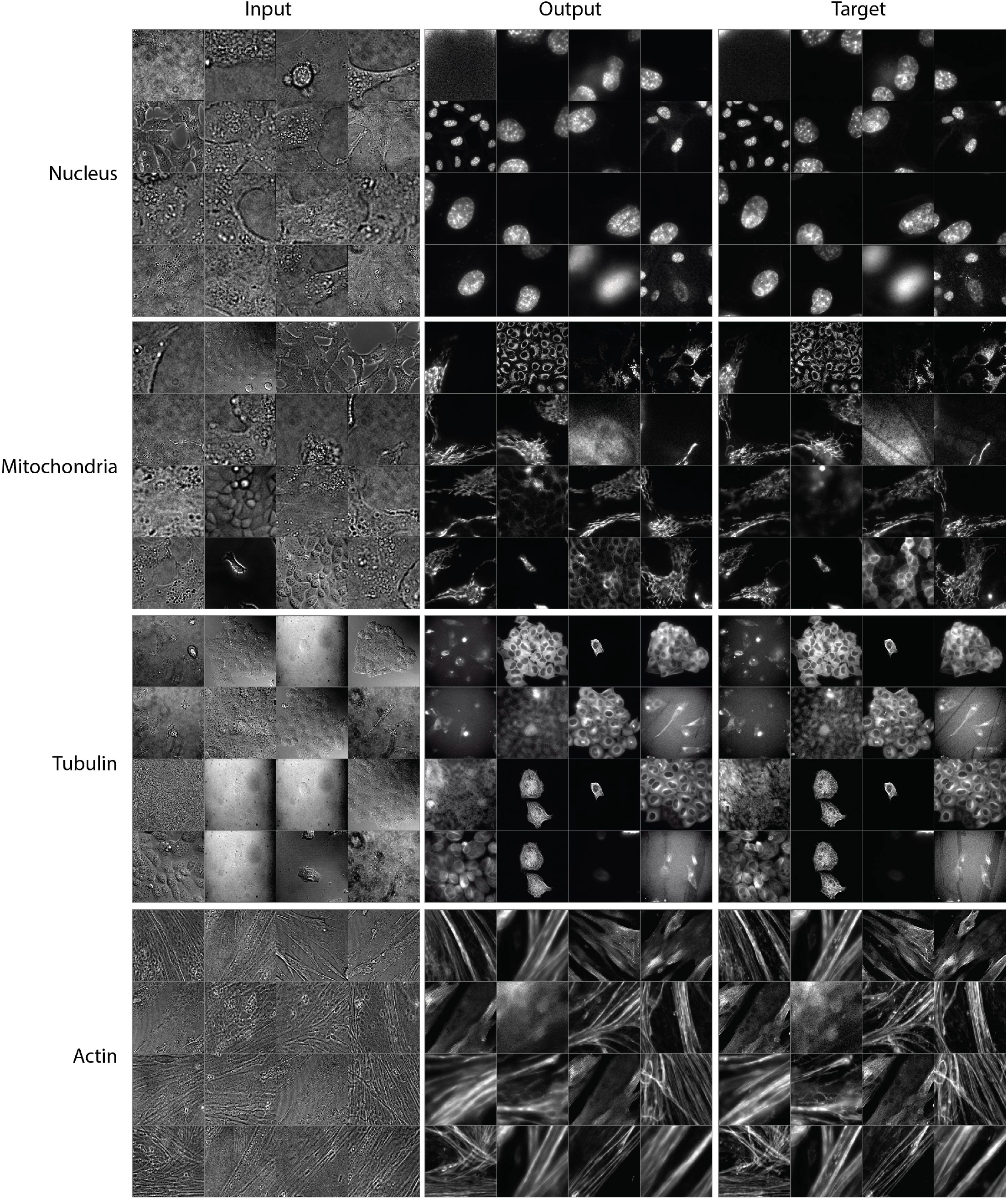
Model outputs on validation images from LMC challenge dataset, sampled at random.

